# Effects of practice on a mechanical horse with an online feedback on performing a sitting postural coordination

**DOI:** 10.1101/2020.07.02.184234

**Authors:** Héloïse Baillet, David Leroy, Eric Vérin, Claire Delpouve, Jérémie Boulanger, Nicolas Benguigui, John Komar, Régis Thouvarecq

## Abstract

The present research aims at quantifying the impact of practicing a new coordination pattern with an online visual feedback on the postural coordination performed on a mechanical horse. Forty-four voluntary participants were recruited in this study. They were randomly assigned to four practice groups based on i) with or without feedback (*i.e*., group 1, control, did not receive the feedback; group 2, 3 and 4 received an online feedback during practice) and ii) the specific trunk/horse coordination to target during practice (group 1, target coordination = 180° (without feedback); group 2, target coordination = 0°; group 3, target coordination = 90°; group 4, target coordination = 180°). All participants performed pre-, practice, post- and retention sessions. The pre-, post- and retention sessions consisted of four trials, with one trial corresponding to one specific target coordination to maintain between their own oscillations and the horse oscillations (spontaneous, 0°, 90°, and 180°). The practice phase was composed of three different sessions during which participants received an online feedback about the coordination between their own oscillations and the horse oscillations.

Results showed a significant change with practice in the trunk/horse coordination patterns which persisted even after one month (retention-test). However, all the groups did not show the same nature of change, evidenced by a high postural variability during post-test for 0° and 90° target coordination groups, in opposition to the 180° and spontaneous groups who showed a decrease in coordination variability for the 180° group. The coordination in anti-phase was characterized as spontaneously adopted by participants on the mechanical horse, explaining the ease of performing this coordination (compared to the 0° and 90° target coordination). The effect of online visual feedback appeared not only on the coordination pattern itself, but most importantly on its variability during practice, including concerning initially stable coordination patterns.

## 1. Introduction

Motor coordination was defined by Bernstein [1] as the process of mastering redundant degrees of freedom involved in a particular movement to make a controllable system. According to the dynamical systems approach of motor control, a coordination corresponds to a coupling mode between different elements of the sensory-motor system [2], emerging from interacting constraints: organism, environmental and task [3,4]. Indeed, Newell [3] defined the constraints as “boundaries of features that limit motion of the entity under consideration” (p.347), therefore reducing the number of possible configurations of a system. Within the coordination dynamics of human motor control literature, posture is a particular case of coordination, viewed by Bernstein [1] as a necessary component for any voluntary motor action. The redundant degrees of freedom of the postural system provide an adaptive means for maintaining balance under the effect of a variety of interacting constraints [3,4]. In most studies, the relative phase (RP) between angular movements of two non-homologue joints, the hip and the ankle, appears to be a natural candidate for describing postural coordination [5–7].

Several studies focusing on target tracking tasks highlighted the existence of two spontaneous coordination pattern: the *in-phase* pattern (*i.e*., RP ≃ 20°) and the *anti-phase* pattern (*i.e*., RP ≃ 180°) [5,8]. The in-phase pattern is characterized by simultaneous flexion or extension of ankles and hips and is exhibited during low oscillation frequencies and small oscillation amplitudes. Conversely the anti-phase pattern is preferred during high oscillation frequencies and large oscillation amplitudes and corresponds to the flexion of one joint associated with the extension of the other joint. These two preferred coordination patterns do not show the same intrinsic stability and previous findings have demonstrated that the stability of the *in-phase* pattern is lower than the stability of the *anti-phase* pattern [6], advocating for pattern stability to be a key feature of postural control.

The variability observed in postural control is known for reflecting the high level of complexity of the postural system, self-organized under many interacting constraints [9]. Hence, several studies have manipulated these constraints, such as in target tracking tasks [5] to determine their effects on the human postural coordination and its stability. Indeed, the influence of the task (*e.g*., keeping a constant distance between the target and the head, tracking the target movement without head movement) had a paramount effect on the stability of the coordination patterns exhibited to maintain balance [7]. The postural coordination can therefore be constrained by properties of the supra-postural task.

In other experiments, participants were instructed to perform a wide range of postural coordination patterns between the hip and ankle (between 0° and 360° for a total of 16 different coordination patterns) [10]. The originality of these studies was based on the addition of behavioral information (Lissajous figure) indicating the coordination mode to adopt. This method was previously used with success in bimanual coordination experimentations [11–13]. Indeed, the Lissajous plot integrates the movement of the two joints into a single point by having the movement of one joint moving the cursor horizontally while the motion of the other joint moves the cursor vertically [11], and by allowing participants to receive supplementary information like online visual feedback to characterize their own postural activity. In these experiments, participants have to reproduce a coordination pattern projected on a screen in front of them with online visual feedback or with a feedback after each trial. Such information allowed participants to produce other coordination patterns in addition to the stable *in-phase* and *anti-phase* patterns [10]. In this case, the use of feedback required an additional intentional charge to the participants that can be characterized as a supplementary task constraint. Indeed, this behavioral information constrained the participant’s intention and the coordination pattern [14].

Another methodology was used by Faugloire [10,15] whereby the participants were informed in real time about the discrepancy between the actual pattern performed and a requested pattern, using two electro-goniometers placed on the hip and on the ankle of participant. Angular movements of the hip and ankle were represented in an orthonormal system with x-axis and y-axis, respectively [10].

Furthermore, the same authors used this method to evaluate the effect of practice on new coordination patterns which can emerge during a supra-postural task [16]. The use of feedback allowed participants to adopt a new postural coordination pattern as well as to modify their initial *in-phase* and *anti-phase* coordination stability [16].

It is to be noted that only a small number of research focused on the study of posture in sitting position although this is a common and familiar position which is actually mastered before the standing posture in human development [17,18]. Most of these studies were therefore performed among infants but also among participants with disabilities [19–22]. Indeed, motor dysfunctions of persons with disabilities may lead to postural problems that results in them spending more time in sitting positions rather than in standing positions to perform vital tasks of daily life [21]. In the sitting position, static and dynamic stability are two important aspects that directly affect body motion or sway [17]. Trunk stability relies on sensory-motor adequacy of body attitude and on adequate muscular responses, constantly modified by interaction of constraints applied on the system. In non-standing positions, postural muscles are active in a cranio-caudal order, with the neck muscles recruited before the trunk muscles [22–24]. Movements of the head are pertinent for exploration of the environment through visual and vestibular systems [17]. According to the same authors, the pelvis can be compared to a rigid body moving around a medio-lateral axis and is therefore a stable support for the trunk. These studies showed the importance of head, trunk and pelvis movements in sitting posture. Hence, compared to Faugloire’s research, the sitting posture cannot be characterized by ankle-hip coordination but instead by the coordination between head, trunk and hip [25,26]. Following this previous study by Faugloire and Baillet [16,27] recently analyzed the sitting coordination according to different angles and showed the key role of the trunk angle. Therefore, the coordination between the trunk and the horse was used as an order parameter in the present study (following [28]). Sitting mostly occurs on static supports like chairs, but it can also occur on dynamical supports like a horse. The latter provides an interesting situation in which the rider has to coordinate cyclical movements to the movement of the horse [29]. To improve and/or study the posture of riders, the mechanical horse was created in the 1990s [30]. The mechanical horse can also be used in rehabilitation centers to improve the motor abilities, muscle tonus, postural coordination and/or energy expenditure of disabled patients [31]. The mechanical horse oscillates in the antero-posterior plane, which affords for major postural coordination modifications in patients [32].

In a previous study, [27] observed that participants unfamiliar with riding adopted a spontaneous pattern of coordination between their trunk and the horse, corresponding to an in *anti-phase* pattern (*i.e*., RP_*trunk-horse*_ ≃ 180°). This pattern was maintained when the horses oscillated at a low frequency whereas an instability and/or a change of coordination pattern appeared when oscillation frequencies increased. This research sparked questions regarding the effects of practice in such a task.

The objective of this present study was to quantify the changes in spontaneous sitting coordination of healthy participants induced from practice on a moving mechanical horse with the assistance of online visual feedback. To assess the intrinsic coordination dynamics, scanning trials (spontaneous, 0°, 90° and 180°; see methodology of [33] for example) were performed, before *(i.e*., pre-test), after *(i.e*., post-test) and one month after the practice period *(i.e*., retention-test). We hypothesized that the practice of a new coordination pattern will modify the spontaneous postural repertoire of participants towards the target pattern. Secondly, we hypothesized that the addition of visual feedback presented online during practice will foster the potential changes in the initial coordination pattern.

## Materials and methods

### 2.1. Participants

Forty-four voluntary participants were recruited for this study. All participants were novice horse riders (*e.g*., no previous experience on real horse neither on mechanical horse), aged between 18 and 30 years old. None of the participants had a history of physical disability or balance disorders. These forty-four participants were randomly assigned into four groups (11 participants in each group). Each group corresponded to the practice of a specific coordination pattern (see Fig. 1): group 1 which was the control group was asked to perform a coordination pattern of 180° between the trunk and the mechanical horse, without receiving any visual feedback; group 2 was asked to perform a coordination pattern of 0° and received online visual feedback during the practice; group 3 was asked to perform a coordination pattern of 90° and received online visual feedback during practice, group 4 was asked to perform a coordination pattern of 180° and received online visual feedback.

**Fig. 1.**
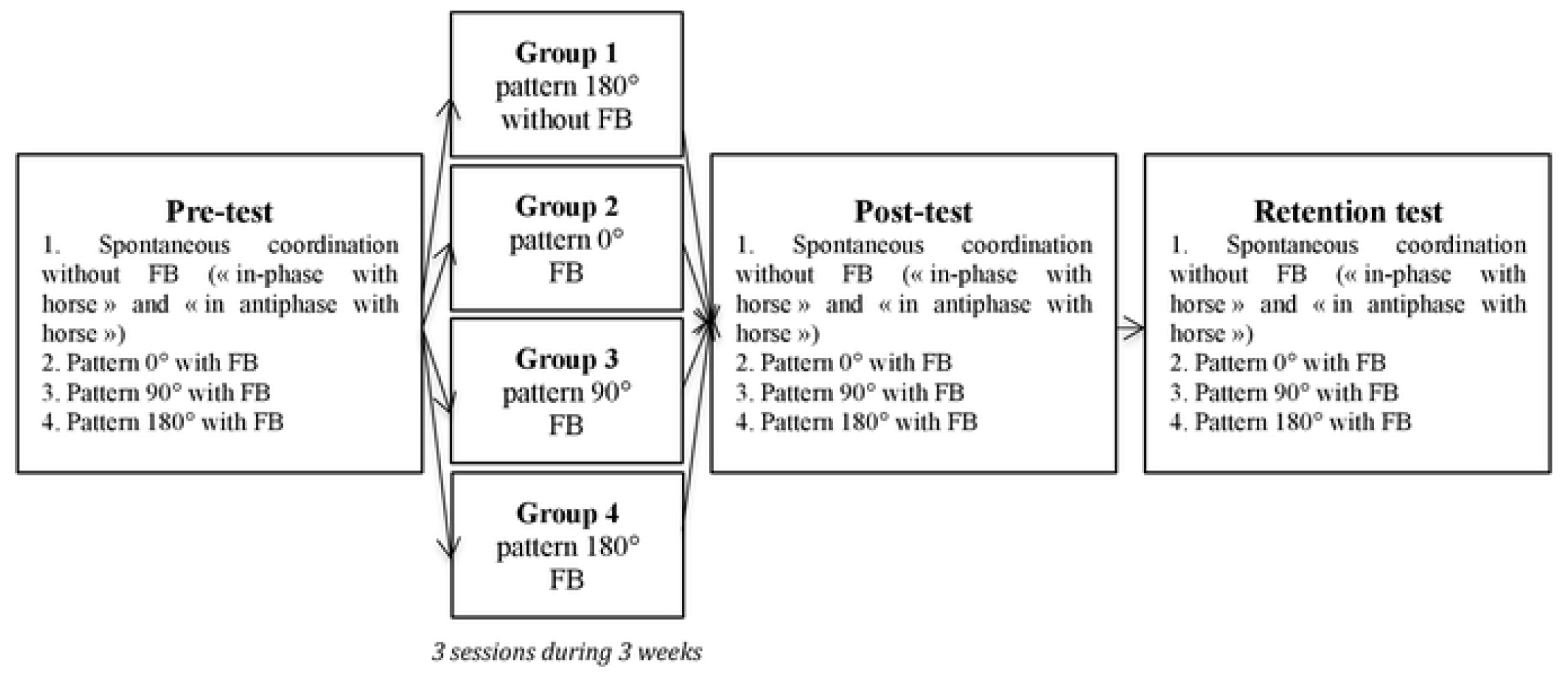
Protocol of study (FB corresponds to Feedback).

### 2.2. Ethics statement

After understanding the purpose of the study, all participants gave written informed consent to participate in the study which is in accordance with the Declaration of Helsinki. The protocol was approved by the human research ethics committee of Lille University (n° 2016-1-S39).

### 2.3. Experimental conditions

The mechanical horse used in this study generated two movements: forward/backward and upward/downward. The amplitude of forward/backward motion was 0.006 meters, or approximatively an amplitude angle of 10°, and the amplitude of upward/downward motion was 0.11 meters. The length of the mechanical horse was 1.74 meters and the oscillation frequency of this tool was adjustable and ranged from 0.2 to 2.5 Hz.

Participants were seated comfortably on the saddle of mechanical horse, their hands holding the reins. Angular positions of participants’ trunk and horse were recorded using two electro-goniometers (Biometrics, Ldt) connected to a DataLog unit (Biometrics, Inc., Gwent, UK). Firstly, to measure trunk oscillations, an electro-goniometer was positioned on the trunk at the level of the right iliac crest of participants, and on the thigh at the level of the greater trochanter (trunk-thigh angle). Secondly, to measure horse oscillations, an electro-goniometer was positioned on the movable part of the mechanical horse and on the fixed part of horse (horse-ground angle), allowing for characterization of the horse oscillations in reference to the horizontal axis. The angular amplitude of this horse was 10°. Each electro-goniometer was sampled at 50 Hz. To estimate the coordination between trunk oscillations and horse oscillations (RP_*trunk-horse*_), the computation of the point-estimate of relative phase (*i.e*., Discrete Relative Phase) using the peak flexion of trunk angle and horse angle [33].

During the experiment, the participants were placed on the mechanical horse that was 2.62 meters away from a screen (2.36 m width x 1.54 m height). The task was to reproduce different postural coordination patterns, 180°, 0°, 90° and 180° according to the allocated group task *(i.e*., control, 0°, 90° and 180° group respectively). The choice of these specific coordination was derived from the study by [32] on brain-damaged patients, in which it was observed that the patients’ spontaneous coordination was close to 0° or 90° contrary to healthy subjects. It was thus necessary to observe the changes (or absence of changes) of these different coordination patterns. The target coordination was visually represented on the screen during 30 seconds at the beginning each trial (Fig. 2). The screen displayed a trunk-horse position plane for which the abscissa and the ordinate axes corresponded to the oscillations of the trunk and to the oscillations of horse respectively, both normalized in amplitude between 1 and -1. The prescribed pattern was presented by a green dot, which was moving on the graph following the requested coordination (Lissajous figure). The 0° pattern corresponded to an oblique line with a positive slope; the 90° pattern corresponded to a circle and the 180° pattern to an oblique line with negative slope (see Fig. 2). On this unique screen, after the initial presentation, the prescribed pattern disappeared and online visual feedback was provided (for 0°, 90° or 180° groups) which corresponded to the actual postural activity of the participant through live streaming of data from the electro-goniometers (*i.e*., trunk and horse oscillations). To give feedback to the participants, the dot turned green when the coordination was close to the target coordination; and it turned blue when the coordination moved away from the target coordination (*i.e*., ± 30° from target coordination).

**Fig. 2.**
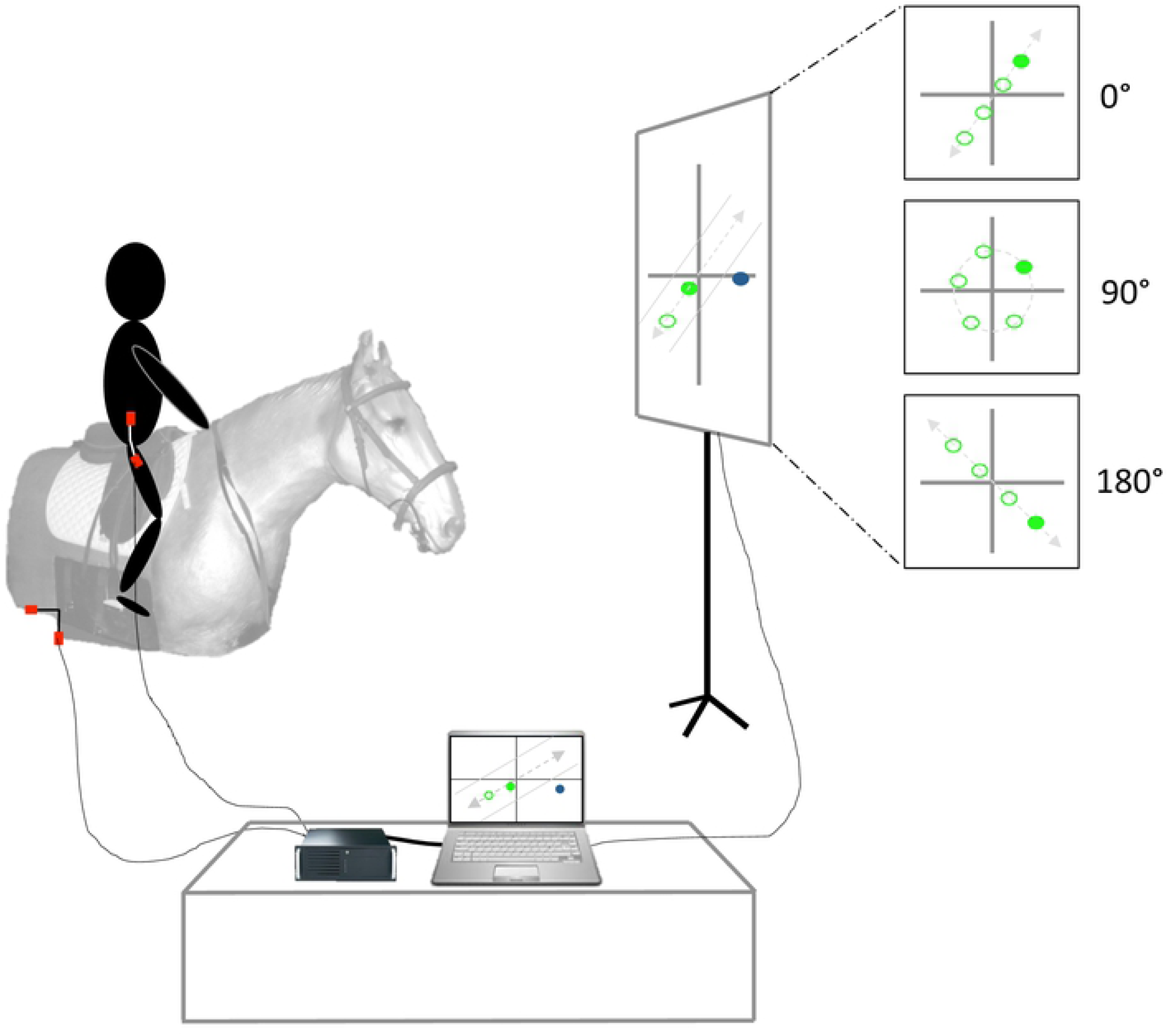
Experimental set-up. Each participant performed 4 different tasks on the mechanical horse; the first coordination was a 180° coordination without feedback and the 3 others were 0°, 90° and 180° coordination with a visual feedback, corresponding to the online postural activity of participant.

The instruction for the participants was to produce trunk oscillations movements (*i.e*., flexion-extension) in accordance with the mechanical horse in order to move the dot on the screen to correspond to the initial observed movement and to keep it green. Participants were also asked to keep seated on the saddle.

The four groups of practice corresponded to a specific target coordination pattern:

- Group 1 (or control group; n=11): the participants were asked to perform a target coordination pattern in *anti-phase* (*i.e*., 180° ± 30°) between the trunk and the mechanical horse, without the help of feedback.
- Group 2 (or group 0°; n=11): the participants were asked to perform a target coordination pattern *in-phase* (*i.e*., 0° ± 30°) between the trunk and the mechanical horse, with the help of online visual feedback.
- Group 3 (or group 90°; n=11): the participants were asked to perform a target coordination pattern in *90° out-of-phase* (*i.e*., 90° ± 30°) between the trunk and the mechanical horse, with the help of online visual feedback.
- Group 4 (or group 180°; n=11): the participants were asked to perform a target coordination pattern in *anti-phase* (*i.e*., 180° ± 30°) between the trunk and the mechanical horse, with the help of online visual feedback.

### 2.4. Procedure

This present study consisted of a pre-test, three practice sessions, a post-test and a retention test (Fig. 1). All participants from each group performed pre-, post- and retention-tests, each test consisting of four trials. Each trial required the participants to perform a different coordination pattern, during three oscillation frequencies of the mechanical horse (chosen through [27]’s study; V1: 0.96 Hz or 50% of maximal horse oscillation frequency; V2: 1.47 Hz or 70%; V3: 1.72 Hz or 80%) each maintained for 3 minutes, with a total of 36 minutes.

The first trial of pre-test (like post- and retention-tests) corresponded to the recording of the spontaneous coordination of each participant. The instruction was “to follow the horse’s movements at each oscillation frequency”, without any indication of results or any real-time visual feedback. Indeed, as with [34]’s study and his protocol with fingers’ coordination, the aim for the participants was to coordinate their movements with that of the horse without instructions on how to coordinate. During the second, third and fourth trials, participants were asked to perform 0°, 90° and 180° target coordination patterns between trunk and horse respectively. During those trials, each coordination pattern was represented on the first screen (in blue) and on the second screen the online visual feedback (in green), giving the participants information on their postural activity in real-time. The instruction provided to the participants was “to get as close as you can to the figure viewed on the first screen; where a green dot, which corresponds to your trunk oscillation, indicates good performance with reference to the target coordination”.

The practice sessions started one week after the pre-test and occurred once a week for three weeks. A practice session consisted of one trial performed for each of the 3 different oscillation frequencies V1, V2, and V3, each maintained for 3 minutes (total of 9 minutes). This trial required a coordination pattern according to the groups: 0° for group 2, 90° for group 3, 180° for group 4 and 180° for control group. For these trials, the same instructions as the one mentioned above were given.

After the 3 weeks of practice, a post-test was performed in a similar sequence as the pre-test. Thereafter, long-term effects were analyzed through a retention test identical to the previous pre- and post-tests, conducted one month after the post-test.

### 2.5. Dependent and Independent Variables

#### 2.5.1. Dependent Variables

Dependent variables were coordinative variables, corresponding to RP_*trunk-horse*_, and its variability (*i.e*., variance of the relative phase, σ^2^, expressed in degrees^2^ and used as a measure of stability [35]), and on the absolute error of target coordination.

#### 2.5.2. Independent Variables

Independent variables were the variables Group, Tests, Target Pattern and Oscillation frequencies. Finally, an independent variables Session was also considered to investigate the evolution of the dependent variables between the 3 practice sessions.

### 2.6. Statistical analysis

Statistical analysis was performed with SPSS software (SPSS Statistics 21, SPSS Inc., IBM, Chicago, IL, USA). The RP_*trunk/horse*_ was considered as circular data (*i.e*., 0°-360°), where 0° and 360° represented the same orientation and the same polar angle. We therefore needed to use circular statistics [36], but circular statistics do not allow the computation of interactions between factors. To perform linear statistics, the range of RP values was thus decreased to be included into the range [0°; 180°]. Indeed, when the range of distribution of values is less than 180°, the difference between circular and linear methods is negligible [15,16,37]. The range was also reduced in [38]: all RP values higher than 180° were subtracted from 360°.

Firstly, to evaluate the effect of feedback on the postural coordination, a statistical analysis was conducted on the value and of RP_*trunk/horse*_ and on the absolute error of target coordination, with a four-way mixed model ANOVA: Groups_(Control/0°/90°/180°)_ * Tests_(Pre-test/Post-test/Retention-test)_ * Target Pattern _(spon/0°/90°/180°)_ * Frequencies_(50%/70%/80%)_, with Tests, Target Pattern and Frequencies as repeated factors. Secondly, the same ANOVA was performed on the variance of RP_*trunk/horse*_ to investigate the effect of practice on individuals’ postural coordination variability. Eventually, four two-way ANOVAs (*i.e*., one for each group) (Sessions_(S1/S2/S3)_ * Frequencies_(50%/70%/80%)_) with both factors as repeated measures were performed to investigate the evolution of the postural coordination (and its variability) across the practice sessions.

For every analysis, the statistical threshold was established at p=.05. When the Mauchly test for sphericity was significant, the Greenhouse-Geisser correction was applied when epsilon was lower than 0.75, otherwise the Hyun-Feld procedure was used. When a significant main effect or interaction was found, the Bonferroni method was used for all post hoc comparisons.

## 3. Results

### 3.1. Trunk/horse coordination

#### RP_*Trunk-Horse*_

The statistical analysis performed on RP_*Trunk-Horse*_ revealed a significant main effect of test F_(1.0,47.7)_ = 6.6 (p=.013), target pattern F_(1.2,49.1)_ = 27.1 (p<.001), and group F_(3,40)_ = 3.2 (p=.033; *d=.19*). Furthermore, the analysis also showed interaction effects between target pattern * group F_(3.7,49.1)_ = 3.9 (p=.009) and test * target pattern F_(1.2,48.0)_ = 5.6 (p=.017).

#### Absolute error of target coordination (Fig. 3)

Regarding the absolute error of the target coordination, the statistical analysis revealed a significant main effect of test F_(2,80)_ = 24.5 (p<.001), target pattern F_(1.5,60.8)_ = 286.1 (p<.001), frequency F_(1.6,63.8)_ = 9.9 (p<.001) and group F_(3,40)_ = 4.6 (p=.007; d=.26). The ANOVA also showed interaction effects between test * group F_(6,80)_ = 2.7 (p=.004), target pattern * group F_(4.6,60.8)_ = 3.2 (p=.015), test * target pattern F_(3.3,133.8)_ = 16.2 (p=.000), target patter * frequency F_(2.8,110.6)_ = 11.6 (p<.001) and test * target pattern * group F_(10.0,133.8)_ = 2.9 (p=.003).

**Fig. 3.**
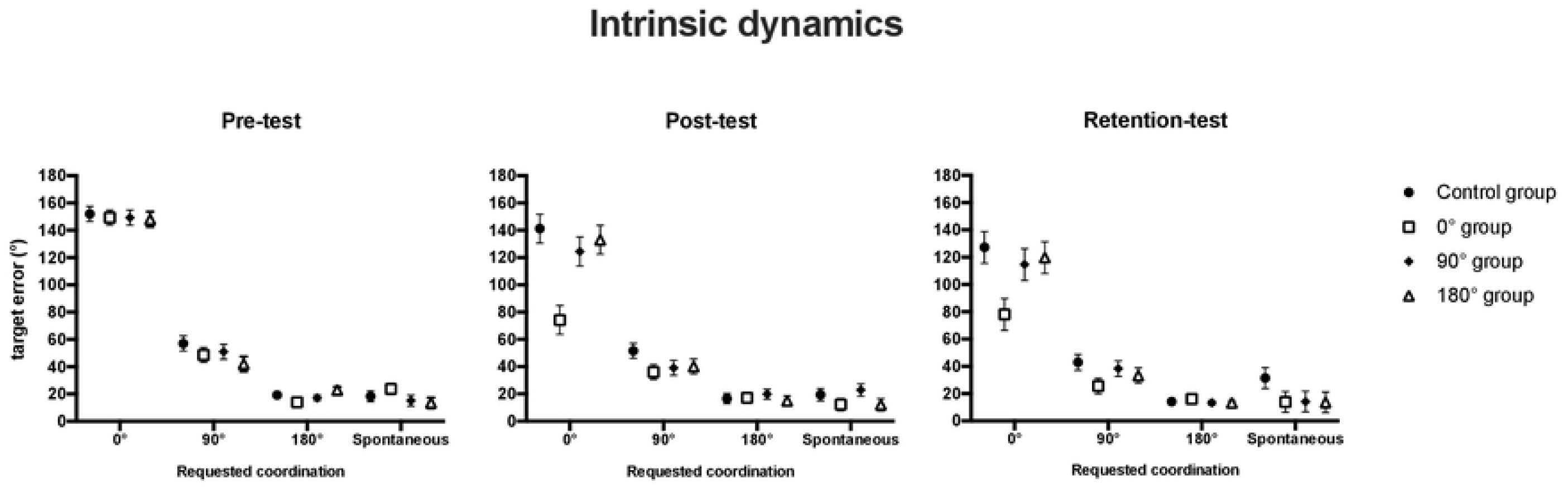
Depictions of coordination intrinsic dynamics: absolute target error of trunk/horse RP (mean ± standard error) for each group at each requested coordination patterns during pre- (left panel), post- (middle panel) and retention tests (right panel).

#### 3.1.1. Interaction between *test* and target coordination

The main results of the post-hoc tests indicated differences in coordination for all groups between pre- (163.0°±7.2°; target error 58.8°±1.3°), post- (140.5°±2.4°; target error 48.5°±1.9°) and retention-tests (143.5°±2.3°; target error 44.3°±2.1°), showing a change of coordination after practice which lasted during the retention phase. The statistical analysis showed that the coordination performed in post-test was different from the ones in pre-test only for the 0° target pattern condition (174.4°±21.8° in pre-test and 118.5°±5.3° in post-test). Similarly, the coordination performed during the retention-test was different from the ones in pre-test only for the 0° target pattern condition (174.4°±21.8° in pre-test and 109.0°±5.8° during retention-test) and the 90° target pattern condition (147.0°±7.8° in pre-test and 120.3°±3.3° during retention-test). More precisely, the statistical analysis performed on absolute target error showed that the coordination error in pre-test was different from the ones in post- and retention-tests for the 0° target pattern condition (149.7°±2.8° in pre-, 118.2°±5.3° in post- and 109.9°±5.8° in retention-tests) and for the 90° target pattern condition (49.6°±2.8° in pre-, 41.9°±2.8° in post- and 35.0°±2.8° in retention-tests). These observations highlighted a coordination change between pre-, post- and retention-tests for the 0° and 90° target patterns, initially close to *anti-phase* during pre-test and shifting closer to the targeted pattern after practice and decreasing the error target. Regarding the spontaneous and the 180° target patterns, the lack of evolution with practice shows that those patterns were already present in the intrinsic dynamics of the participants.

#### 3.1.2. Interaction between group and target pattern

Regarding the group effect, it appeared that the significant difference occurred between the 0° practice condition (137.3°±8.0°; target error 42.4°±5.3°) and 180° practice condition (158.4°±8.0°; target error 50.*5°*±*3.5°)*. However, it seems that the exhibited coordination was different according to group but also according to the target pattern. Indeed, for the control group, the actual coordination measured when the target pattern was 90° (139.5°±6.6°) was different compared to the 180° target pattern (163.3°±2.1°) and to the spontaneous target patterns (162.9°±2.3°); for the 0° group, the coordination performed during the 0° target pattern (100.5°±13.8°) and the 90° target pattern (115.2°±6.6°) was different from the coordination exhibited during the 180° target pattern (164.3°±2.1°) and during the spontaneous target pattern (169.2°±2.3°); for the 90° group, the coordination during the 90° target pattern (137.3°±6.6°) was different from the coordination exhibited during the 180° target pattern (164.8°±2.1°) and during the spontaneous target pattern (163.1°±2.3°); for the 180° group, the coordination performed during the 90° target pattern (137.1°±6.6°) was different from the three other target patterns (166.4°±13.8° in 0° target pattern, 163.2°±2.1° in the 180° target pattern and 167.1°±2.3° in spontaneous target pattern). More precisely, the values of absolute target error when the target pattern was 0° (error of 140.1°±8.1° for control group, 100.5°±8.1° for 0° group, 129.4°±8.1° for 90° group and 133.5°±8.1° for 180° group) were different compared to those of 90° (50.6°±4.4° for control group, 36.8°±4.4° for 0° group, 43.0°±4.4° for 90° group and 38.3°±4.4° for 180° group), those of 180° (16.7°±1.9° for control group, 15.7°±1.9° for 0° group, 16.7°±1.9° for 90° group and 17.0°±1.9° for 180° group) and those of spontaneous target patterns (23.1°±3.2° for control group, 16.6°±3.2° for 0° group, 17.4°±3.2° for 90° group and 13.1°±3.2° for 180° group).

#### 3.1.3. Interaction between group, test and target pattern

The statistical analysis performed on target error showed significant differences between tests as a function of groups and target pattern. The post-hoc tests revealed no significant differences of coordination error for the control group in pre- (61.7°±2.5°), post- (57.2°±3.8°) and retention-tests (53.9°±4.3°). However, the target error measure during pre-test for 0° was different from the post- and retention-tests. On the other hand, for the 90° and 180° groups these differences were found between pre- and retention-tests. Moreover, it appears that these coordination errors were different as a function of the predicted target pattern. During post- and retention-tests, for control, 0°, 90° and 180° groups, measured coordination errors measured during 0° and 90° target patterns were different from the three other target patterns. These observations highlighted the decreasing of the absolute target error after practice sessions for the 0°, 90° and 180° groups.

### 3.2. Variability of trunk/horse coordination (Fig. 4)

The statistical analysis performed on the variability of RP_*Trunk-Horse*_ revealed significant effect of prescribed target pattern F_(1.8,73.2)_ = 84.9 (p<.001), oscillation frequency F_(2,39)_ = 19.7 (p=.000), but no effect of group (p=.523; *d=.05*). This analysis also showed interaction effects between *test* * target pattern F_(3.8,152.1)_ = 5.8 (p<.001), target pattern * frequency F_(6,35)_ = 5.3 (p=.001), target pattern * frequency * group F_(12.7,169.4)_ = 1.9 (p=.036), and *test* * pattern * frequency F_(6.7,267.5)_ = 2.2 (p=.037).

**Fig. 4.**
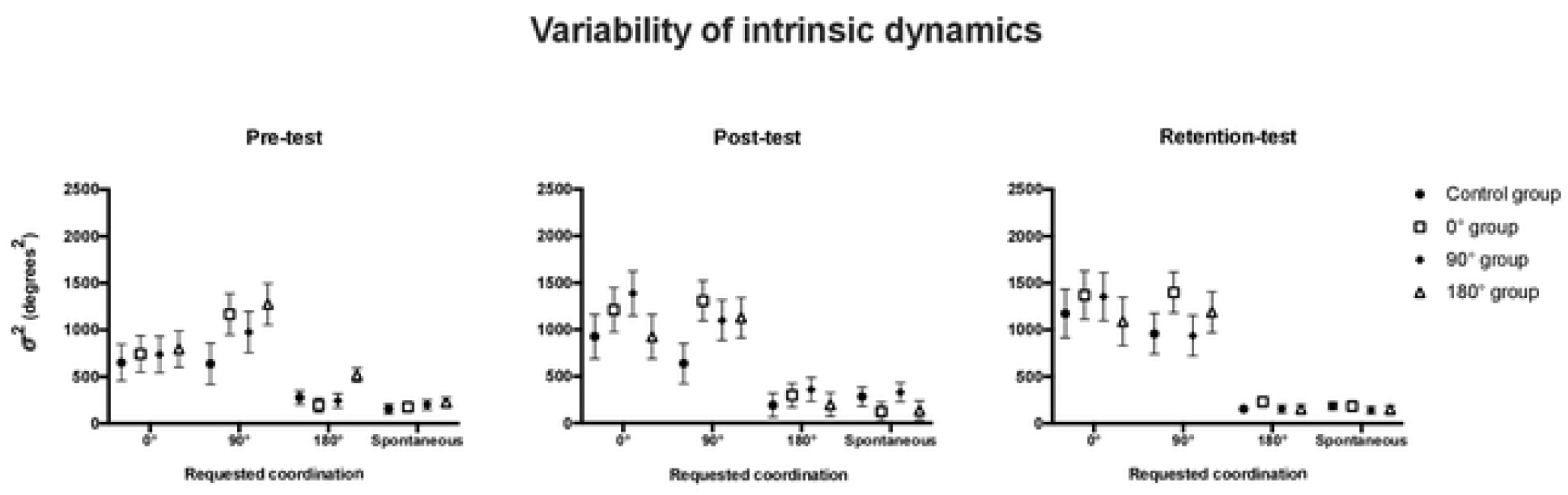
Depictions of the variability of coordination intrinsic dynamics: variability (σ^2^, in degrees^2^) of trunk/horse coordination (mean ± standard error) for each group at each requested coordination patterns during pre- (left panel), post- (middle panel) and retention tests (right panel).

Focusing on the first three ways interaction, *i.e*. the pattern * frequency * group effect, significant differences between groups were observed only at the 90° target pattern and at maximal frequency. Indeed, the postural variability measured for the 0° group (1376.0°±173.3°) was different from the variability exhibited by the control group (571.6°±173.3°) and from the variability exhibited by the 90° group (601.4°±173.3°), showing a higher stability for the participants in the control and the 90° groups compared to the variability of the participants in the 0° group.

In the same vein looking at the second three ways interaction, *i.e*. the test * target pattern * frequency effect results showed that the postural variability performed during the 0° target pattern in pre-test (868.1°±143.9°) was different from the variability during post-test (1390.9±150.8°) and retention-tests (1621.3°±162.3°) in the 50% horse oscillation frequency condition, as well as in the 70% horse oscillation frequency condition (631.7°±119.1° in pre-test, 1167.1°±157.8° in post-test, 1219.7°±163.0° in retention-test). Furthermore, for the 180° target pattern, at 50% horse oscillation frequency, the variability during pre-test (418.3°±56.8°) was different from the variability during retention-test (198.8°±18.4°). These results show clearly an increase in postural variability for the 0° target pattern across the sessions. However, results show a decrease in this variability for the 180° target pattern. It should be noted, however, that the increase in variability was less pronounced when the oscillation frequency was higher.

### 3.3. Evolution of trunk/horse coordination

#### 3.3.1. Comparison between four groups in pre-test

A comparison was performed between the four groups (control, 0°, 90° and 180° groups) at pre-test, for the coordination and variability of coordination. Therefore, a statistical analysis was performed on the trunk/horse coordination and on variability of RP_Trunk-Horse._ These comparisons showed no group effect at pre-test regarding the relative phase (F_(3, 40)_=0.81; p=0.49) and variability (F_(3, 40)_=1.12; p=0.35). The participants’ behavior (and the behavior variability) of each group was similar before the practice session. Evolution of trunk/horse coordination and their variability as a function of practice sessions for each group (Fig. 5):

**Fig. 5.**
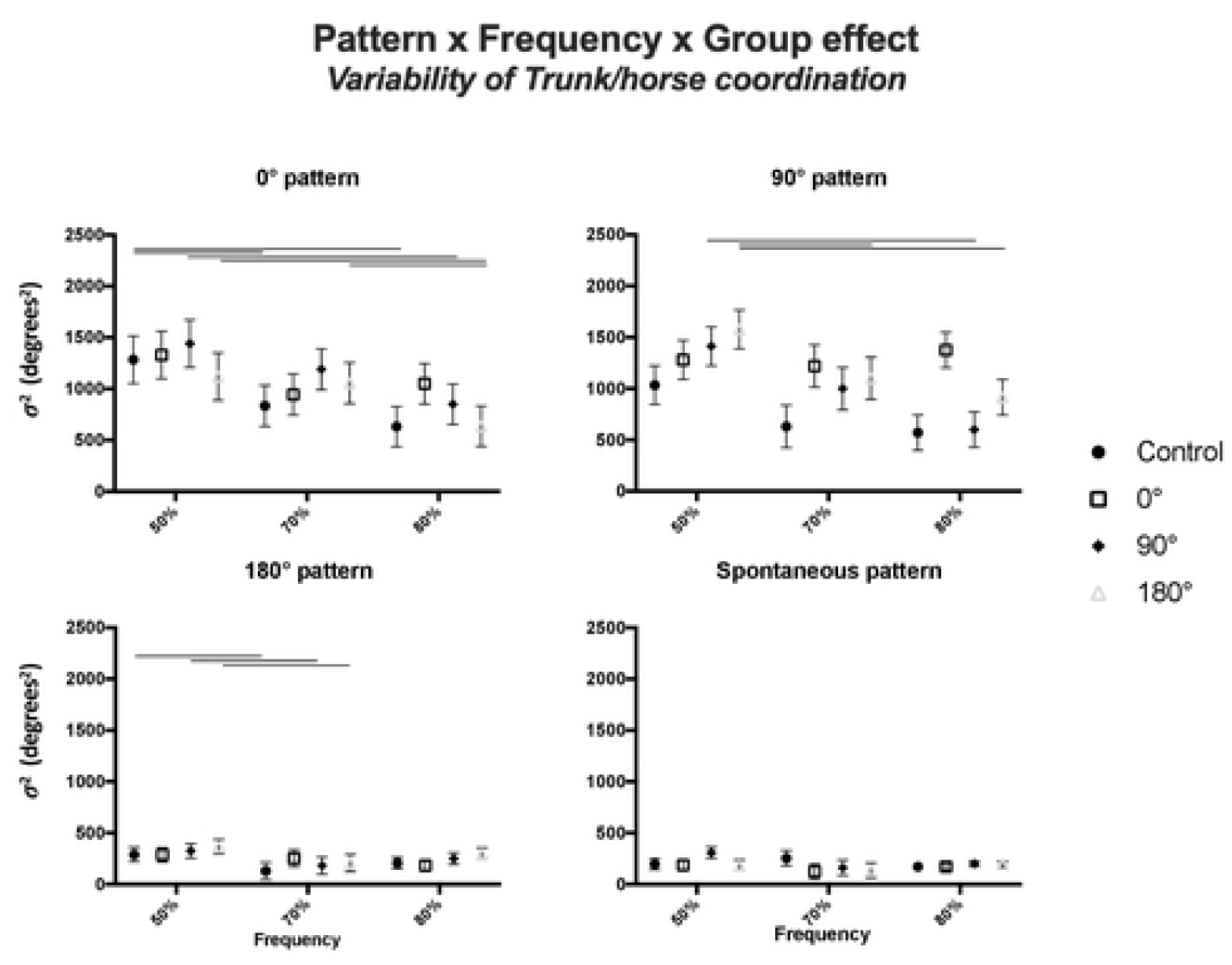
Depictions of the relative phase (left panel) and variability (right panel) of trunk/horse coordination (mean ± standard error) measured for all groups at each practice sessions (S1, S2 and S3). The horizontal lines characterize significant differences between sessions.

#### 3.3.1. Control group

For both the trunk-horse coordination value and for its variability, the two-way ANOVA performed on the sessions for control group revealed no significant effect and no interaction effect between the sessions and frequencies. Results therefore showed no effect of practicing the intrinsic coordination pattern without any feedback.

#### 3.3.2. Group 0° target pattern

For the group who practiced the 0° coordination, a significant effect of the practice session was shown F_(2,9)_ = 11.1 (p=.004), but no effect was found for horse oscillation frequency and no interaction effect was evidenced (p>.05). *Post-hoc* tests showed that the coordination of 0° group measured at the first session (133.9°±8.3°) was different from the two other sessions (104.3°±12.2° at S2 and 91.9°±11.0° at S3). Moreover, for the postural variability, a significant effect of practice session was shown for the 0° group, F_(2,9)_ = 6.5 (p=.018). The postural variability decreased between session 2 (828.1°±220.9°) to session 3 (185.6°±24.4°).

#### 3.3.3. Group 90° target pattern

The two-way ANOVA showed a significant effect of the practice session F_(2,9)_ = 4.73 (p=.039), and a significant effect of horse oscillation frequency F_(1.3,12.5)_ = 5.5 (p=.048), but no interaction effect was evident. Regarding the session effect, Bonferroni post-hoc tests showed a unique difference between the coordination at session 1 and 2 (133.5°±5.9° at S1, 117.2°±9.5° at S2 and 122.8°±5.7° at S3). Moreover, this test performed on effect of horse oscillation frequency showed that the coordination at 50% frequency (108.4°±7.4°) was different from the coordination performed at 70% (131.7°±8.0°). The statistical analysis performed on the postural variability of the 90° group indicated a significant main effect of the practice session F_(2,9)_ = 4.4 (p=.047), but no interaction effect between sessions and frequencies. *Post-hoc* tests primarily showed that the variability during session 1 (894.5°±198.5°) was different from the variability during session 3 (243.6°±59.1°). The values of these variances indicate a decrease in the postural variability during practice sessions for this group who practiced the 90° target pattern.

#### 3.3.4. Group 180° target pattern

No significant change in the trunk/horse coordination had been shown (p>.05) throughout the sessions. However, when the statistical analysis was performed on the postural variability, a significant effect of practice session F_(2,9)_ = 8.2 (p=.009) and an interaction effect sessions * frequencies F_(4,7)_ = 4.4 (p=.043) appeared. More precisely, the significant interaction effect followed by post-hoc comparisons showed a main trend of decrease in coordination variability; however this decrease in variability was not identical across the different horse oscillation frequencies. Indeed, in the 50% horse oscillation frequency condition, the postural variability was lower only during session 3 (*i.e*., showing a significant difference only between variability during session 2 (1194.3°±326.6°) and session 3 (339.6°±11.6°)). Conversely during the 80% horse oscillation frequency condition, the variability during session 2 (633°±192°) and session 3 (294.7°±121.7°) was lower than the variability during session 1 (1756.1°±252.6°), when no difference appeared during session 2 and session 3. In other words, if the practice tended to decrease the coordination variability, this decrease appeared later at a higher horse oscillation frequency.

## Discussion

The aim of this study was to quantify the impact of practice on a new coordination pattern, with the help of live visual feedback, on the spontaneous postural coordination performed by healthy participants sitting on an oscillating mechanical horse.

### 4.1 Effect of practice on the mechanical horse on participants’ initial repertoire

Validating our hypothesis, the results showed a significant modification of trunk/horse coordination with practice, after only three practice sessions of a specific pattern. More precisely, the postural coordination measured through scanning trials during the post- and retention-tests was different from the coordination during the pre-test. A statistical analysis stated that all participants presented a similar variability and coordination in pre-test, confirming these modifications. These modifications of postural coordination with practice have been already observed in several studies performed on the bimanual coordination [39] or on the postural coordination (hip/ankle coordination in standing posture) [40].

Furthermore, with the present results, it seems that the practice of a specific pattern allowed participants to be closer to the novel 0° and 90° target pattern *(e.g*., 174° in pre-test, 118° in post-test, and 109° in retention-test), suggesting the possible capacity of participants to perform a new coordination that was not initially present in their repertoire before practicing. This may be confirmed by increasing the number of sessions.

Observing the postural coordination, absolute target error and variability performed by all participants during all the different target patterns, it appears that the 0° and 90° target patterns were the most impactful. Indeed, the values of target error (Fig. 3) during 0° and 90° target patterns were greater and the variability values (Fig. 4) indicated an important variability of the coordination during those patterns. Whereas during the 180° and spontaneous target patterns, the initial behavior was less impacted regarding its stability and possibility in moving closer to the target. More precisely, the postural variability measured in post- and retention-tests did not show a decrease compared to pre-test for all groups, with sometimes an increase in this variability during post- or retention-tests (Fig. 4). However, the target error values decreased between pre-, post- and retention-tests. The literature specifies that the learning of a new coordination pattern is often synonymous with postural destabilization [15,41–43]. Actually, an increase in variability associated with a decrease of the in-target error can be the sign of an early stage of learning, when increased variability can help to leave an initial coordination pattern therefore playing a functional role in the learning process. Previous studies from [44–46] showed that movement variability can be beneficial for learning but can appear with different delays between participants (*e.g*., appeared only after ten learning sessions [44]). In that sense, the limited number of practice sessions performed by the participants could explain the lack of decrease in variability with practice. Nevertheless, the coordination variability measured was different as a function of the target coordination pattern requested during pre-, post- and retention-tests. Indeed, participants showed high variability and high target error during the 0° and 90° target patterns, with a significant increase of coordination variability *and* a significant decrease of the target error between pre-, post- and retention-tests demonstrated by the 0° practice condition group. On the other side, coordination variability and target error were lower for the 180° target pattern and spontaneous pattern, with a significant decrease between pre- and retention-tests demonstrated by the 180° practice condition group. These results confirmed that participants experienced difficulty when performing the non-spontaneous patterns: 0° and 90°, evidenced by a highest level of variability *and target error* in those conditions. These results could validate that the 0° and 90° practice condition groups were in an early stage of learning, whereas the spontaneous and 180° practice condition groups had already passed this early stage of learning as the coordination patterns practiced in their respective groups were already in their initial repertoire.

Observing the effect of the horse oscillation frequency on individual postural variability, it appears that as the frequency increases, the variability decreases. In other words, the postural stability of participants increases as a function of horse’s oscillations, even during the 0° and 90° target patterns where the global level of coordination variability is higher. However, the coordination values and target error indicated that it seems more difficult for participants to maintain the target coordination pattern when the horse frequency increases. For example, when the 0° pattern must be performed, the mean coordination was close to 123° at low frequency (0.96 Hz), and when the oscillation frequency increased, at medium and high frequencies (1.47 and 1.72 Hz), this coordination approached the *anti-phase, i.e*. 180°. This 180° pattern can be considered as a strong attractor as all the participants tend to fall into this pattern when the control parameter increases [33]. Indeed, all participants seemed to be attracted by the *anti-phase* coordination between trunk and horse when the horse frequency increases [5]. These results show that the oscillation frequency impacts postural coordination, but also its stability. In other words, increasing the level of constraints during practice tends to push every participant towards an identical behavior and to increase the stability of this identical pattern. Practicing at high frequency could then impair the ability to leave the initial attractor by limiting the functional variability from occuring. Those results confirm that of previous studies performed on standing posture [5,6,8,47,48] and on bimanual coordination [49–51]. Indeed, the *anti-phase* coordination observed in existing studies is a stable postural coordination.

According to the specific groups, differences regarding the ease of reproducing the practiced coordination pattern were observed after the three practice sessions. The 0° and 90° practice condition groups seem to be the most affected by practice. Indeed, instead of the control group and the 180° practice condition group who exhibited the required coordination, the 0° and the 90° practice condition group never really reached the target coordination. However, trying to practice a specific coordination, accompanied by a live visual feedback, seemed enough to lead to a change in the initial repertoire of the participants. Indeed, the practice condition only allowed the individuals of these 0° and 90° groups to get closer to the target coordination without actually reaching it. This result confirms that of previous studies, such as [5,6,10,15,16,41,52–57], showing the possibility of influencing the postural repertoire of a participant even if he never really reaches the target coordination during the practice.

### 4.2 Effect of the online visual feedback on practice

Focusing on the practice sessions instead of the scanning trial sessions (remember through statistical analysis, that the behavior of all participants was similar during pre-test), results showed singular evolution in the practiced coordination (and target error) and its variability. Specifically, for the group who practiced the 0° coordination, the participants trunk/horse coordination tended to get closer to the practiced pattern (*e.g*., during session at 70%, 142.8°±28.7°, during session 2 at 110.6°±48.8° and during session 3 at 90.1°±56.6°). On the other end, there were no observed differences between the sessions for the control and 180° practice groups. This result is mainly due to the fact that the practiced coordination was already in the repertoire of the participants and hence, the study did not require them to adopt a new pattern [27]. However, the 180° practice condition group displayed some effect from the practice sessions in terms of the variability of the coordination pattern – which was not exhibited by the control group. In other words, using a live visual feedback can help to temporarily destabilize an existing coordination pattern, whereas the simple practice of a coordination without the feedback definitely has no effect on this coordination. This result goes beyond previous results from [10] that showed no effect of online feedback on practice, as results of the current study demonstrate the potential benefits of online feedback in helping the participant move out of his initial repertoire. Eventually, the addition of an online visual feedback had a larger impact, leading to a shift towards the target coordination by impacting the stability of an already existing pattern (*e.g*., when the target coordination is not in the initial repertoire of the participant).

More precisely, the postural variability analysis during practice sessions shows a reduction of the variance throughout these sessions. For groups with online visual feedback (0°, 90° and 180°), we noticed a significant decrease in this variability between session 1 and session 3. This decrease in variability with practice for the groups receiving an online visual feedback can be explained mainly by a high variability of coordination during the initial sessions and a variability during the last session identical to the control group. This high variability of the coordination for the 0°, 90° and 180° groups during the first sessions can be explained by the decrease in the strength of the attractor, a prerequisite to any potential modification of the attractor landscape [58]. Although the addition of such an additional informational constraint could be viewed as negatively affecting the coordination and therefore the postural stability of the individual [10], observations on the three groups who received the online feedback indicate a potentially beneficial effect of feedback through incorporating functional variability during practice. Those results therefore go beyond previously cited studies of [10,40], which did not distinguish the real benefit of visual feedback in learning a specific coordination pattern, by looking at how online visual feedback can infuse early variability in practice that could later lead to better performance [59].

## Conclusion

Through this study, it has been shown a significant change with practice of new trunk/horse coordination patterns which persisted even after one month, with only three practice sessions given to perform the new coordination patterns. The effect of online visual feedback appeared not only on the coordination pattern itself, but most importantly on the variability of coordination patterns during practice, including initially stable coordination patterns. This method using visual feedback has demonstrated its efficacy for healthy subjects, with the next potential step being to transfer these results for rehabilitation purposes. Specifically, to improve the postural coordination of brain-injured patients and to help them relearn coordination patterns lost as a result of lesions (for example, an *anti-phase* coordination of 180°). Future studies should confirm these results to improve the existing rehabilitation protocols. Nonetheless, some limitations need to be highlighted from the present study, specifically regarding the retention test. Indeed, the retention test was still performed with Lissajous information for the three coordination patterns (0°, 90° and 180°), as the objective was to look at the effect of practicing with the online feedback. In this regard, [60], have noted that participants may become dependent on the feedback when performing bimanual coordination tasks. With the aim of looking at the comparative effect of this online feedback on learning and transfer, it will be interesting to perform all the tests in the exact same condition, i.e. without online feedback (such as the spontaneous coordination condition). Thus, a future study should address this limitation.

## 3. Acknowledgments

The research reported here was supported by the CPER-FEDER Q7 project [Axe: Masses de Données et Contenus Intelligents, project: Data science: méthodologies et applications (DAISI)] and the Institut Français du Cheval et de l’Equitation (IFCE), whom we thank. The authors thank Dr Nadège Rochat for his participation and help during this study.

